# Expression of Bovine Annexin A4 In E. *coli* Rescues Cytokinesis Blocked by Beta-Lactam Antibiotics

**DOI:** 10.1101/2023.07.14.549091

**Authors:** Carl E. Creutz

## Abstract

Treatment of bacteria with beta-lactam antibiotics can impair the process of cytokinesis, the final step in cell division, leading to the formation of a filamentous form of the bacteria. The expression of a mammalian calcium-dependent, membrane-binding protein, bovine annexin A4, in *E. coli* was found to reverse the inhibitory effects on cytokinesis of the beta-lactam antibiotics ampicillin, piperacillin, and cephalexin. This novel activity of the annexin was blocked by mutation of calcium binding sites in the annexin, indicating roles for calcium binding to the annexin and the binding of the annexin to membranes in restoring cytokinesis. The filamentous form of the bacteria has been reported to be more resistant to phagocytosis by cells of the immune system in eukaryotic hosts. Therefore, expression of annexins in pathogenic bacteria, by promoting the breakdown of the bacterial filaments, might serve as an adjuvant to enhance the efficacy of beta-lactam antibiotics.

## INTRODUCTION

*E. coli* and many other bacteria under conditions of stress reduce the frequency of undergoing cytokinesis while continuing to grow and replicate their genome. The result is transformation of the bacteria into a filamentous form[1]. For infectious bacteria this may have survival value by preventing consumption of the bacteria by phagocytic cells of the immune system of the host organism, and the formation of biofilms that protect bacterial colonies. This report describes the unexpected discovery that the expression of mammalian annexin A4, a calcium-dependent, phospholipid-binding protein, in *E. coli* tends to restore the ability for bacteria undergoing antibiotic stress to undergo cytokinesis and leads to a return to a normal morphology.

The annexins constitute a highly conserved family of calcium-binding proteins in plants, animals, and protists [2, 3]. However, they are not expressed in *E. coli*. Motifs similar to the canonical fourfold calcium- and membrane-binding motifs in eukaryotic annexins [4] have been discovered in proteins in some strains of bacteria [5, 6]. Their occurrence in bacteria is rare, however. The bioinformatic analysis of Kodavali et al [6] detected annexin motifs in only 17 bacterial species among the thousands of genomes analyzed. It is unclear if the bacterial annexin-like proteins evolved independently from eukaryotic annexins or if genetic material encoding annexin motifs was acquired by non-genetic transfer from the eukaryotic relatives. The bacterial annexin-like proteins are unusual in that they do not generally follow the pattern of the fourfold domain repeat found in the eukaryotic examples as they have as few as a single annexin motif and up to 6 copies of the repeats. None of the bacterial proteins containing annexin motifs have yet been isolated or characterized biochemically so it is not known if they bind calcium and membranes, although the sequence similarities to the eukaryotic proteins suggest this may be the case. Nor is there evidence for any particular function of the bacterial annexins obtained, for example, by gene deletion or modification experiments. Therefore, the ability of the eukaryotic annexin to modulate a specific function in bacteria, cytokinesis, was not anticipated.

The affinity of eukaryotic annexins for calcium is modulated by the coincident binding of phospholipids [7, 8] so the annexins are implicated in many activities regulated by calcium that occur with or on membranes. It has been reported that inhibition of the expression of a particular eukaryotic annexin, annexin A11, in human A431 and HeLa cells leads to the failure of cytokinesis in its final stage during which the neck of cytoplasm between daughter cells undergoing mitosis collapses and the membranes fuse, separating the cells [9]. It is not known what the precise role for the annexin may be in this process, but regulation by calcium, and the ability to pull two membranes together to promote membrane fusion could provide an explanation consistent with the known in vitro properties of the annexins [10].

In 1940 Gardner reported that, while working with Chain, Florey, and others at Oxford, he observed that many types of bacteria exposed to penicillin at low concentrations formed long filaments [11, 12]. Subsequently it was determined that this morphological change was caused by the binding of penicillin or other beta-lactam antibiotics to proteins that play essential roles in bacterial cell division[13–15]. For example, in E.coli penicillin binds to penicillin binding protein 3 (PB3) which functions in peptidoglycan synthesis in the bacterial cell wall, including in the septa between dividing cells. This process is inhibited by the binding of penicillin resulting in continued growth of the bacteria without cell division, producing filaments containing multiple copies of the bacterial genome.

It has been suggested that the formation of these filaments may provide a selective advantage to bacteria under attack by phagocytic cells of the immune system of eukaryotic organisms which are unable encapsulate the extended bacterial filaments [1, 16]. To this extent the formation of bacterial filaments may be of medical importance in that they may impair treatment of bacterial infections with beta-lactam antibiotics. The result is that bacteria gain resistance to the host defenses precisely when the antibiotic is administered with the intention to inhibit bacterial growth. It is possible that small molecule drugs that could inhibit the filamentation process might provide an effective adjuvant treatment to enhance antibiotic efficacy, but no such compounds have been identified. The results reported here suggest that expression of annexins in pathogenic bacteria might serve such a prophylactic benefit.

## MATERIALS AND METHODS

### Materials

Components of the bacterial growth medium, Luria broth, (10 gm/ liter tryptone, 5 gm/liter NaCl, yeast extract 10 gm/liter) were obtained from Fisher (Fair Lawn, New Jersey). Bacterial agar for petri plates was from Acros Organics (The Hague, Netherlands), ampicillin from Fisher (BP1760), piperacillin from Sigma-Aldrich (P8396) (St. Louis, Missouri), cephalexin from Alpha Aesor (J63172) (Haverhill, Massachusetts), IPTG from Fisher, and glutaraldehyde from Fisher. Stock concentrations of the antibiotics were prepared as follows and stored at minus 20 degrees C: ampicillin 100 mg/ml in 1:1 ethanol:H2O; piperacillin 2 mg/ml in ethanol; and cephalexin 25 mg/ml in 1:1 ethanol:H2O. Other standard chemicals were of reagent grade obtained from Fisher or Sigma.

*E. coli* strain BL21(DE3) was obtained from Thermofisher Scientific (Waltham, Massachusetts) and the expression plasmid pET11d from Millipore Sigma (Burlington, Massachusetts).

Annexin expression plasmids were constructed as described [17, 18]. Since the plasmids were originally sequenced manually in the early 1990’s the inserts in all plasmids used in this study were resequenced with high fidelity automated Sanger sequencing by Azenta Life Sciences, (Burlington, Massachusetts). All designed mutations were confirmed and no spurious mutations were found.

Unique reagents will be provided by the author upon reasonable request.

## Methods

Darkfield light microscopy of bacterial cultures was performed with a Nikon Labophot microscope (Nikon Instruments, Melville, New York) with a dark field condenser (dry) and an e plan40X/0.65 Nikon objective. Micrographs were recorded with an Amscope Microscope digital camera MU 1803-HS 18mp Aptina color CMOS (Irvine, California) using software supplied with the camera.

Cell culture and microscopy. Protein expression from the plasmid pET11d can be regulated by the addition of the inducer IPTG (Isopropyl β-d-1-thiogalactopyranoside) [19]. However, protein expression from this plasmid can have significant baseline expression without the inducer. It was seen in SDS gels of extracts of bacteria expressing annexin A4 that basal expression of the annexin appeared to be 5 to 10% of expression seen with the inducer [20]. In preliminary tests for the present study it was found that including the inducer (concentration 0.4 mM, previously used for maximum protein production) led to formation of inclusion bodies of insoluble protein, distortion of bacterial morphology and partial lysis and aggregation of the bacteria. Accordingly, all of the imaging and experiments described in this report were performed without adding the inducer, which avoided these problems.

Five ml precultures in LB medium with 200 µg/ml ampicillin were seeded with colonies from LB-ampicillin agar plates and incubated on a rotator overnight at 34 degrees C. Fifty µl samples of the saturated cultures were diluted into 5 ml of LB medium with the specified antibiotics in 17X100 mm clear polystyrene culture tubes with loose fitting caps that permitted gas exchange (Fisher Healthcare 14-956-6B) and the growth of the cultures at 34 degrees C was monitored by dark field microscopy and measurement of absorbance at a wavelength of 600 nm. For the microscopy, 20 ul samples of the culture were placed on a glass slide (Fisher, 12-550123 1 × 3 inches, 1 mm thickness, precleaned by the manufacturer) and the drop of culture fluid covered with a cover slip (Fisher 12540B 22X22 mm, 0.17-0.25 mm thickness). Slides were rested for 2 to 5 minutes to allow the cells to settle before observation. Nonetheless, the cells were not tightly adherent to the slides and occasionally some cells moved slightly during imaging resulting in a double image. Cultures were checked and micrographs taken at at 5 to 20 min intervals for up to 4 hours. Occasionally during this period the culture tubes were briefly inserted into a Genesys 20 spectrophotometer (Thermospectronic, Rochester, NY) and the absorbance at 600 nm was recorded to measure culture density. The culture tubes have a circular cross section with an inside diameter of 1.3 cm, so the absorbances recorded were approximately 30% higher than would have been measured in a square cuvette with a one cm pathlength.

For preparation of the micrographs in Figures 1 through 3 comparing the morphology of the different transformed strains, each test culture (5 ml) was inoculated at the same time with 50 ul of the saturated overnight preculture. For display in the Figures, photographs taken after approximately the same period of growth of each of the parallel cultures were obtained when 20 µl samples provided a microscope image containing a broad sampling of the cell morphologies seen without overcrowding. Each experiment illustrated was repeated at least three times with comparable results.

**Figure 1:**
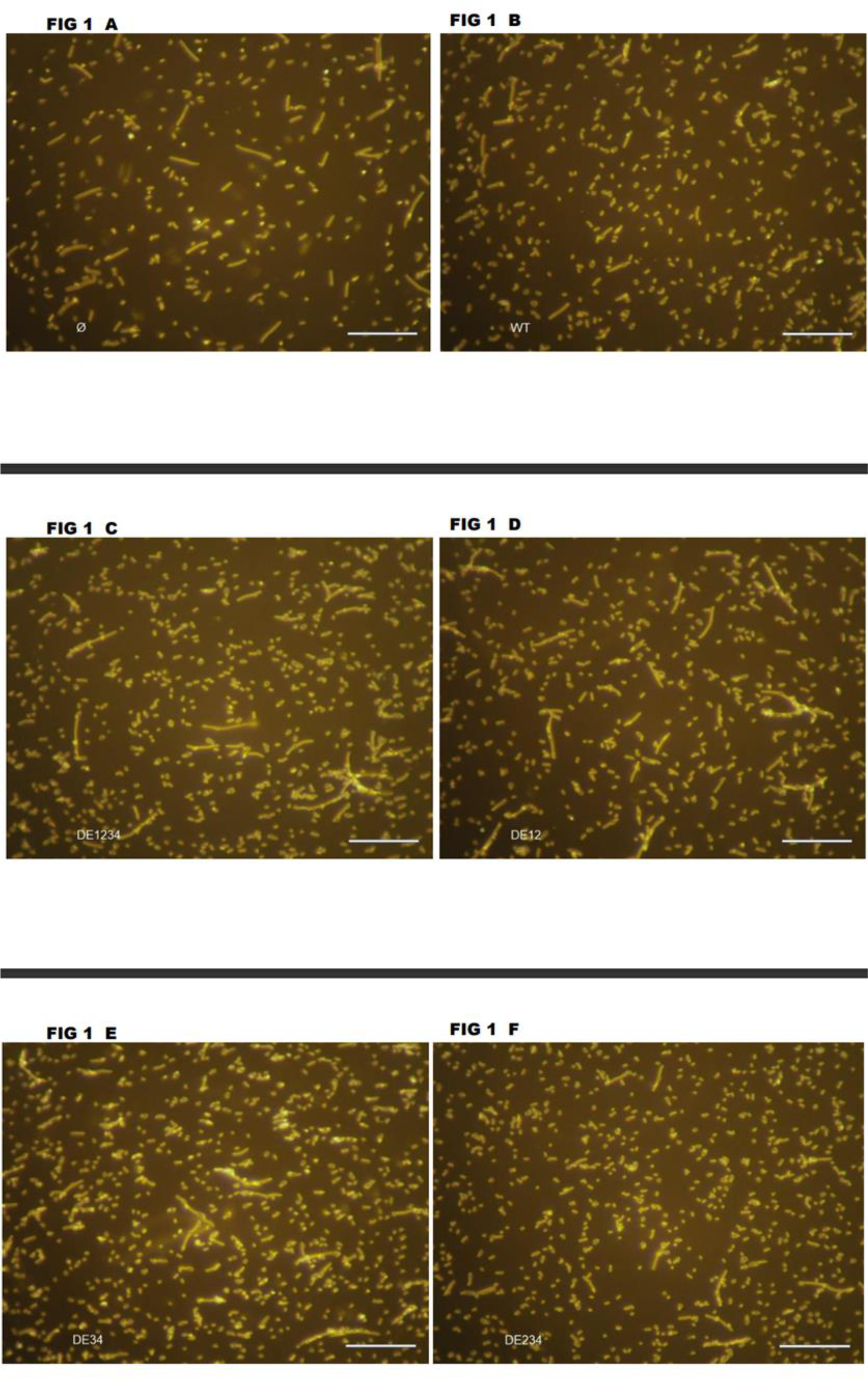
Dark field light micrographs of *E. coli* grown in AMPICILLIN (200 µg/ml) and expressing wild type or mutant annexin A4. **1A:** Ø, control plasmid with no insert **1B**: WT wild type annexin A4 **1C:** DE1234 mutant annexin A4 **1D:** DE12 mutant annexin A4 **1E:** DE34 mutant annexin A4 **1F:** DE234 mutant annexin A4

Electron microscopy*. E. coli* samples selected for ultrastructural analysis were processed for electron microscopy using technical support and instrumentation in the Advanced Microscopy Facility at the the University of Virginia School of Medicine. The samples were fixed in glutaraldehyde and embedded in epon. Ultrathin sections were stained with uranyl acetate and lead citrate and visualized with a JEOL JEM1230 transmission electron microscope.

## RESULTS

### Morphology of E. coli expressing wild type and mutant annexins A4 (the “DE” mutants)

In 1993 Michael Nelson described the construction of a combinatorial library of mutant annexin A4 cDNAs that was designed to facilitate study of the roles of each of the four homologous calcium and membrane binding sites in bovine annexin A4 in biochemical studies[18][17]. The annexin cDNAs in the library were expressed in *E. coli* using the prokaryotic expression vector pET11d. Each calcium binding site contains an important acidic amino acid residue, aspartate or glutamate, that contributes to the coordination of calcium in the cation binding site. Replacement of these residues with alanine by site directed mutation was used to create a library of mutant proteins with reduced calcium and membrane binding properties, which correlated with the specific combinations of mutations present. These mutants were called the “DE” mutants, for replacement of aspartate (D) or glutamate (E). The sites of the mutations are denoted by the domains harboring the mutations. For example, the DE1 mutant has the calcium binding site in the first domain mutated. The DE12 mutant has the calcium binding sites in the first and second domains mutated, and so forth up to the DE1234 mutant which has all four sites mutated. The specific residues mutated in the constructs used in the present study are listed in the paper by Nelson and Creutz [17]. These mutations reduced the calcium sensitivity of the annexin by up to an order of magnitude in two types of assays: binding of the annexin to a biological membrane (chromaffin granule membrane) and aggregation of chromaffin granule membranes. The calcium sensitivities were altered depending upon which mutations, or combination of mutations were introduced (see Nelson and Creutz [17], Table 4). As demonstrated in the sections below, the use of selected mutants from this library revealed that the ability of the annexin to promote cytokinesis in bacteria was dependent on the competence of some of these sites.

### Control cultures grown in ampicillin (200 µg/ml) without expression of the annexin (Fig 1A)

Dark field light microscopy of cultures of *E. coli* strain BL21DE3 harboring the expression plasmid pET11d without an annexin cDNA insert grown in LB-ampicillin media reveals a broad distribution of bacterial cell lengths from 1 to 20 µm.

### Bacterial cultures grown in ampicillin expressing wild type annexin A4 (**Fig 1B**)

When the bacteria were transformed with the expression plasmid containing the wild type annexin A4 cDNA, the number of longer, filamentous forms of the bacteria were reduced relative to the shorter forms, although the optical density (A600) of the cultures with or without the annexin were similar, suggesting net growth of bacterial mass was not greatly altered.

### Bacterial cultures grown in ampicillin expressing the DE1234 mutant annexin A4 (**Fig 1C**)

When bacteria were transformed with the expression plasmid containing the DE1234 mutant annexin, with all four calcium binding sites mutated, a number of longer filaments were formed that were up to 50 µm long – longer than the ones that formed when the bacteria were transformed with the empty expression vector (Fig 1A). It therefore appeared that the annexin in which all the calcium binding sites had been mutated had a slight dominant negative effect on the process of cytokinesis in a subset of cells since longer filaments resulted. However, most of the bacteria remained small, 2 to 5 µm in length.

### Bacterial cultures grown in ampicillin expressing the DE12 mutant annexin A4 (**Fig 1D**)

When bacteria were transformed with a plasmid expressing the DE12 mutant, which is defective in the calcium binding sites in domains 1 and 2, but has normal calcium binding sites in domains 3 and 4, the cultures had a distribution of sizes similar to control cells transformed with the empty plasmid (Fig 1A). This suggests that domains 1 and 2 play a dominant role in mediating the cytokinesis promoting effect, while domains 3 and 4 play a minor role in promoting this effect.

### Bacterial cultures grown in ampicillin expressing the DE34 mutant annexin A4 (**Fig 1E**)

When the bacteria were transformed with the plasmid expressing the DE34 mutant, which has the calcium binding sites in domains 3 and 4 mutated but retains wild type calcium binding sites in domains 1 and 2, there were fewer long filaments and short 1 to 5 µm bacteria were more prominent. The culture therefore resembled cultures transformed with the plasmid expressing the annexin with wild type calcium binding sites in all four domains (Fig 1B). The calcium binding sites in the first two domains of the annexins are the most highly conserved in sequence and the results suggest these two sites provide the major contribution to the cytokinesis promoting activities of the unmutated protein.

### Bacterial cultures grown in ampicillin expressing the DE234 mutant annexin A4 (**Fig 1F**)

When bacteria were transformed with the construct expressing the DE234 mutant the culture looked most similar to cultures expressing the wild type annexin (Fig 1B). This result is notable in that this mutant only has one functional calcium-binding domain – domain 1 – so this single domain appears to be sufficient to support the cytokinesis promoting activity of the intact annexin.

### Studies with ampicillin (200 µg/ml) plus piperacillin (4 µg/ml) (**Fig 2**)

Ampicillin binds penicillin binding protein 3 (PB3)[15]. A related beta-lactam antibiotic, piperacillin, binds PB3 with higher specificity for PB3 than other beta-lactam antibiotics and at low concentrations affects the formation of septa between dividing bacterial cells to a greater extent than elongation of cells [21]. Piperacillin was tested in this study as it was thought it might be more effective at promoting filament formation by the bacteria and therefore provide a better model for detecting the effects of the annexin on cytokinesis. As described in the sections below, differences between the DE mutants in affecting cytokinesis were much more apparent in cultures treated with ampicillin plus piperacillin than in cultures described above that were treated with ampicillin alone.

**Figure 2:**
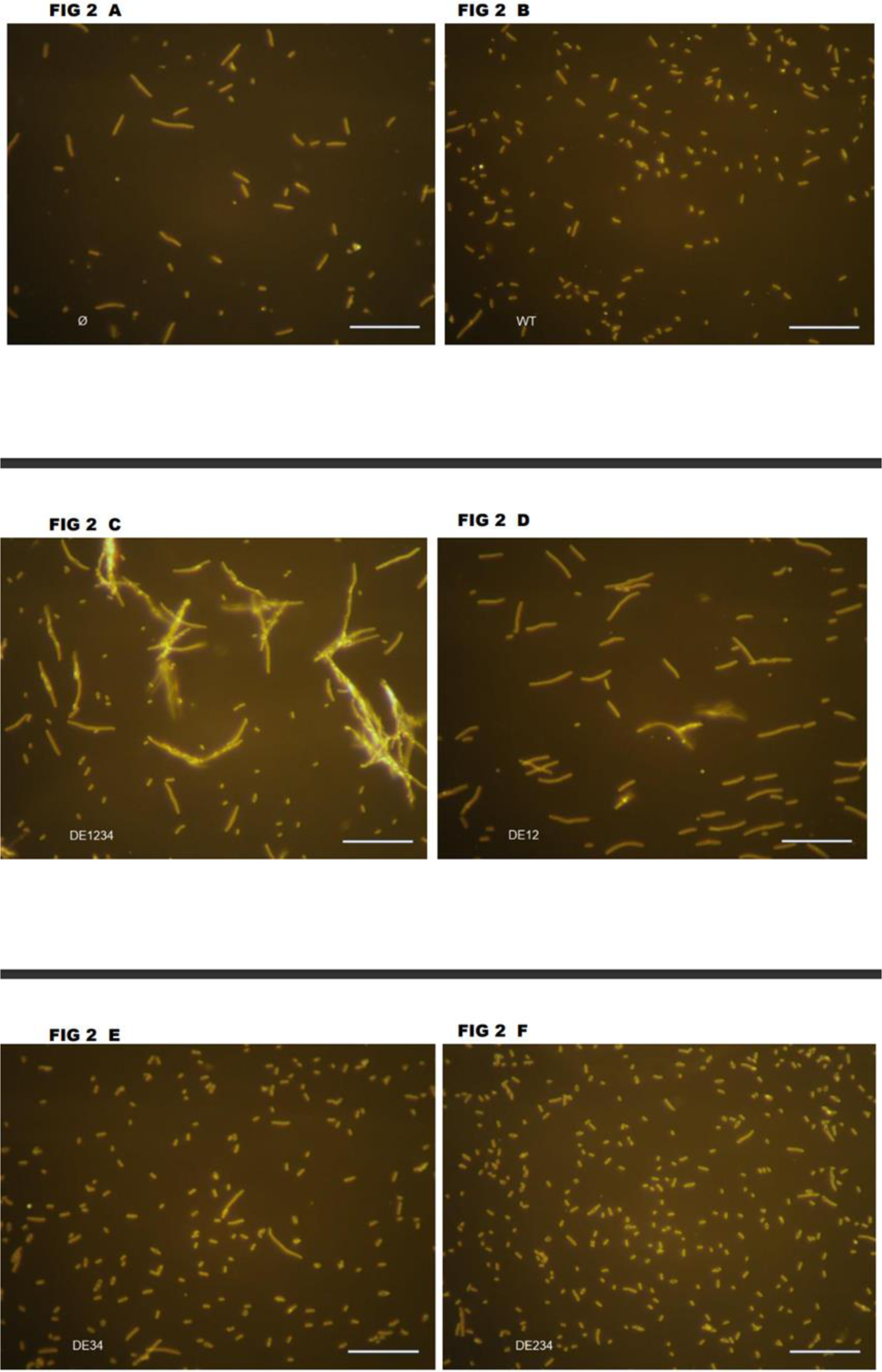
Dark field light micrographs of *E. coli* grown in AMPICILLIN (200 µg/ml) PLUS PIPERACILLIN (4 µg/ml) and expressing wild type or mutant annexin A4. **2A:** Ø, control plasmid with no insert **2B**: WT wild type annexin A4 **2C:** DE1234 mutant annexin A4 **2D:** DE12 mutant annexin A4 **2E:** DE34 mutant annexin A4 **2F:** DE234 mutant annexin A4

### Control bacterial cultures in ampicillin plus piperacillin without expression of annexin A4 (Fig 2A)

As shown in Fig 2A, bacteria harboring the empty expression plasmid and grown in 200 µg/ml ampicillin supplemented with 4 µg/ml piperacillin predominantly formed intermediate length filaments 10 to 40 µm long, with very few bacteria of near normal 2 to 4 µm length.

### Bacterial cultures in ampicillin and piperacillin expressing wild type annexin A4 (Fig 2B)

When the bacteria were transformed with the plasmid expressing the wild-type annexin A4 (Fig 2B) the filament lengths were predominantly reduced to a length of 2 to 10 µm.

### Bacterial cultures in ampicillin and piperacillin expressing the DE1234 mutant annexin A4 (Fig 2C)

When the bacteria were transformed with the plasmid expressing the DE1234 mutant (Fig 2C), the cells formed longer filaments than were seen with the plasmid without an insert, suggesting the DE1234 quadruple mutant had a dominant negative effect on cytokinesis as was also seen above when the cells were cultured in ampicillin alone (Fig 1C). In addition, the longer filaments tended to aggregate, which may have been due to partial lysis of the cells (Fig 2C).

### Bacterial cultures in ampicillin and piperacillin expressing the DE12 mutant annexin A4 (Fig 2D)

When the bacteria were transformed with the plasmid expressing the DE12 mutant in medium containing ampicillin plus piperacillin, the filaments were significantly longer than normal, typically 10 to 40 µm (Fig 2D). Therefore, the loss of the first two calcium binding sites apparently significantly impaired the ability of the annexin to promote cytokinesis in the presence of piperacillin .

### Bacterial cultures in ampicillin and piperacillin expressing the DE34 mutant annexin A4 (Fig 2E)

When the bacteria were transformed with the expression vector for the DE34 mutant the predominant length of the filaments was reduced to 5 to 10 µm in piperacillin plus ampicillin, indicating, as before with ampicillin alone, the mutants with intact first and second calcium binding domains alone were more effective at promoting cytokinesis than mutants that had only the intact third and fourth calcium binding domains.

### Bacterial cultures in ampicillin and piperacillin expressing the DE234 mutant annexin A4 (Fig 2F)

When the bacteria were transformed with the plasmid expressing the DE234 mutant, the cells formed predominantly short filaments of 2 to 8 µm length in ampicillin plus piperacillin, indicating once again, that the presence of only a single functional calcium binding site in domain 1 was sufficient to promote cytokinesis comparable to, or slightly greater than the extent that was promoted by the wild type annexin.

### Studies with ampicillin (200 µg/ml) plus cephalexin (25 µg/ml) (Fig 3)

The antibiotic cephalexin has one of the highest affinities among the beta-lactam antibiotics for binding to penicillin binding protein 3 [15] and was found in this study to have the greatest propensity to promote the formation of long, stable filaments, compared with ampicillin and piperacillin. When bacteria transformed with the empty expression plasmid (**Fig 3A**), or with the wild type annexin (**Fig 3B**), or the DE1234 mutant (**Fig 3D**) were grown in ampicillin supplemented with cephalexin, long filamentous cells were formed over time, suggesting cytokinesis was blocked nearly completely. Many of the filaments extended beyond the field of view in the micrographs and the lengths were not measured. However, the filaments formed with the DE234 mutant (**Fig 3C**) were of limited length at the same time point (3 hours and 20 minutes after seeding the cultures) and were estimated to be 15 to at least 100 µm in length. These filaments were observed over time to be growing more slowly than the other constructs, apparently because both growth and cytokinesis were occurring simultaneously.

**Figure 3:**
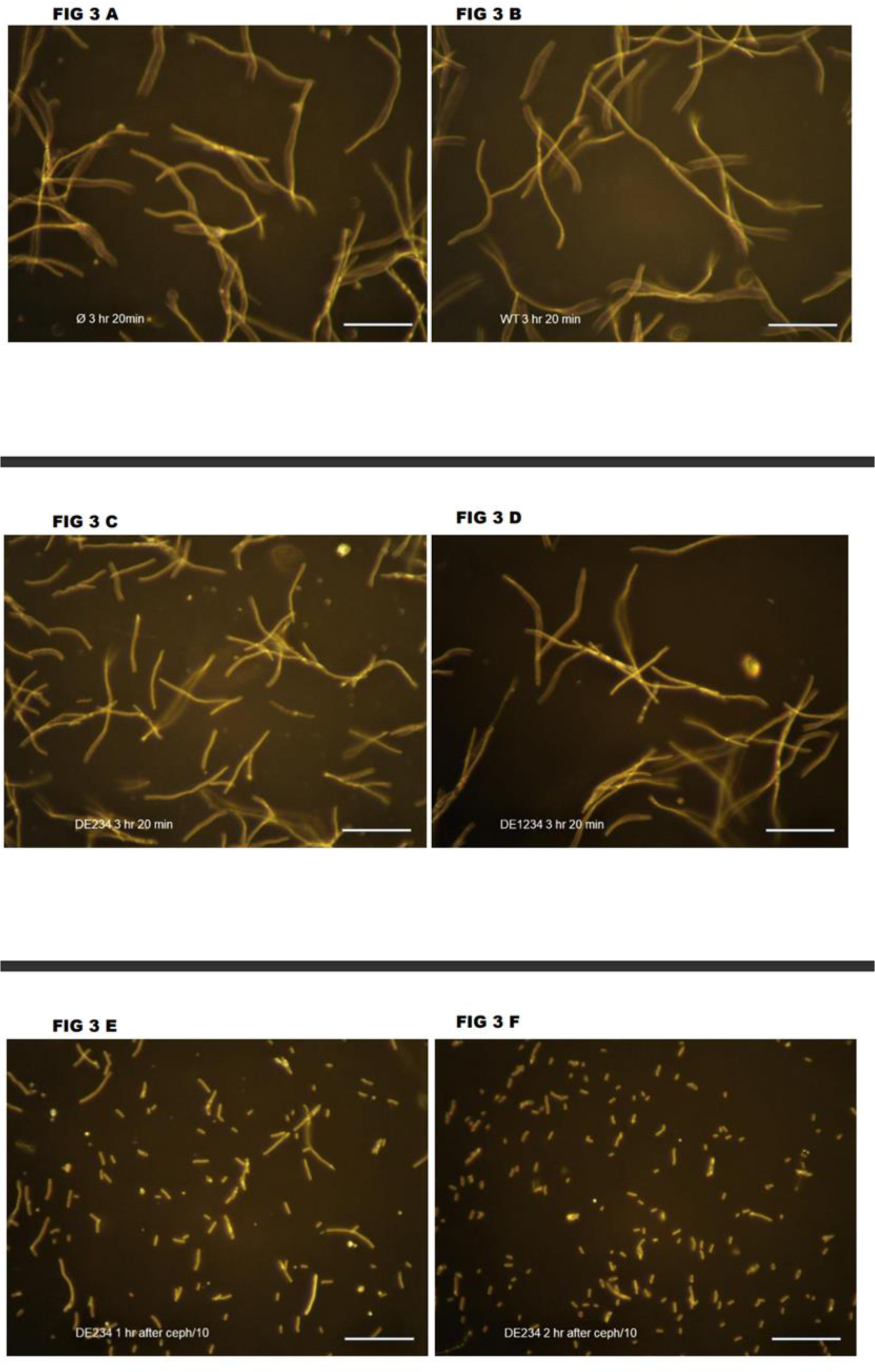
Dark field light micrographs of *E. coli* grown in AMPICILLIN (200µg/ml) PLUS CEPHALEXIN (25 µg/ml) and expressing wild type or mutant annexin A4. **3A:** Ø control plasmid with no insert; 3 hr 20 min, time after initial inoculation of the culture **3B**: WT wild type annexin A4; 3 hr 20 min, time after initial inoculation of the culture **3C:** DE234 mutant annexin A4; 3 hr 20 min, time after initial inoculation of the culture **3D:** DE1234 mutant annexin A4; 3 hr 20 min, time after initial inoculation of the culture **3E:** DE234 mutant annexin A4; 1 hr after ceph/10 (this is the culture from Fig 3C after diluting the cephalexin 10-fold and continuing the incubation for 1 hr) **3F:** DE234 mutant annexin A4; 2 hr after ceph/10 (this is the diluted culture from Fig 3E after waiting another hour during which cytokinesis continues – see text for additional details).

Although the length of the filaments of the DE234 mutant cells grew more slowly than the lengths of the other constructs, in all cases the optical densities of the cultures increased at approximately the same rate, plus or minus approximately 20%. Apparently the mass of total bacterial cells in the culture was increasing at a similar rate independent of the presence of the mutations, but the mass was differentially distributed between cells in the extended, filamentous form and the more normal shape as shorter rods, without having a large effect on the relationship between optical density and mass of cell material present.

### Tenfold dilution of the cephalexin concentration results in an increase in cytokinesis in the filaments formed with the DE234 mutant, but not in the filaments formed with other mutants (Fig 3E and 3F)

After 3 hours and 20 minutes of growth in the presence of ampicillin and cephalexin the cultures were diluted tenfold in fresh medium containing ampicillin but no cephalexin, resulting in a cephalexin concentration reduced from 25 to 2.5 µg/ml to test the stability of the filaments. The filaments formed by bacterial cultures harboring the empty plasmid, wild type annexin, or the DE1234 quadruple mutant were completely stable for the next two hours, unaffected by the reduction in cephalexin concentration (not shown). However, the cells expressing the DE234 mutant, containing only one functional calcium binding site, began to undergo cytokinesis to form shorter filaments after one hour, and further divided by two hours after the dilution. (Compare **Fig 3C** with **3E** and **3F**). This activity of the DE234 mutant parallels the superior ability of the same mutant to promote cytokinesis in ampicillin alone or in ampicillin plus piperacillin seen in **Figures 1** and **2** above

### Electron microscopy of cells expressing wild type or mutant annexins A4

Cultures of the BL21(DE3) cells harboring the empty expression plasmid or plasmids expressing wild-type or selected mutant annexins A4 were grown in LB medium plus 200µg/ml ampicillin and 4 µg/ml piperacillin and processed for electron microscopy of ultrathin sections of pelleted cells to determine if the expression of the annexin altered the ultrastructure of the cells, particularly at sites of cell division. No unique morphology was apparent at sites of cell division with or without annexin expression. There was no apparent increase of protein density at division sites as might be expected if the annexin were preferentially accumulated at sites of cell division and membrane fusion. However, in all of these experiments the expression level of the annexin was necessarily kept low since high levels of annexin expression resulted in disruption of membrane structure and aggregation of the bacteria, as described in the Methods section.

Although no unique ultrastructure was seen in cells expressing the annexin, the changes in filament length due to the annexin, as seen by light microscopy, were confirmed in the electron micrographs. **Fig 4A** shows the morphology of cells harboring the empty expression plasmid and therefore without any annexin. This representative figure shows that there are both short bacterial cells present and filamentous forms, as was seen in the light micrographs of these cultures. **Fig 4B** shows cells expressing the wild type annexin A4. In this case the number of short cells exceeds the number of extended filamentous forms due to the cytokinesis enhancing activity of the annexin. Similarly, in **Fig 4C** the number of short cells exceeds the filamentous forms due to the cytokinesis promoting activity of the DE234 mutant, while in **Fig 4D** the filamentous forms exceed the short forms due to the enhancement of filamentous forms due to the action of the DE1234 mutant. In these electron micrographs of ultrathin sections it is not possible to determine if what appear to be short cells in the electron micrographs are not actually cross sections of the filamentous forms. However, light micrographs of similar cultures (e.g. in **Fig 2**) show the complete cells and support the relative abundance of short and long forms in the cultures as interpreted in the electron micrographs.

**Figure 4:**
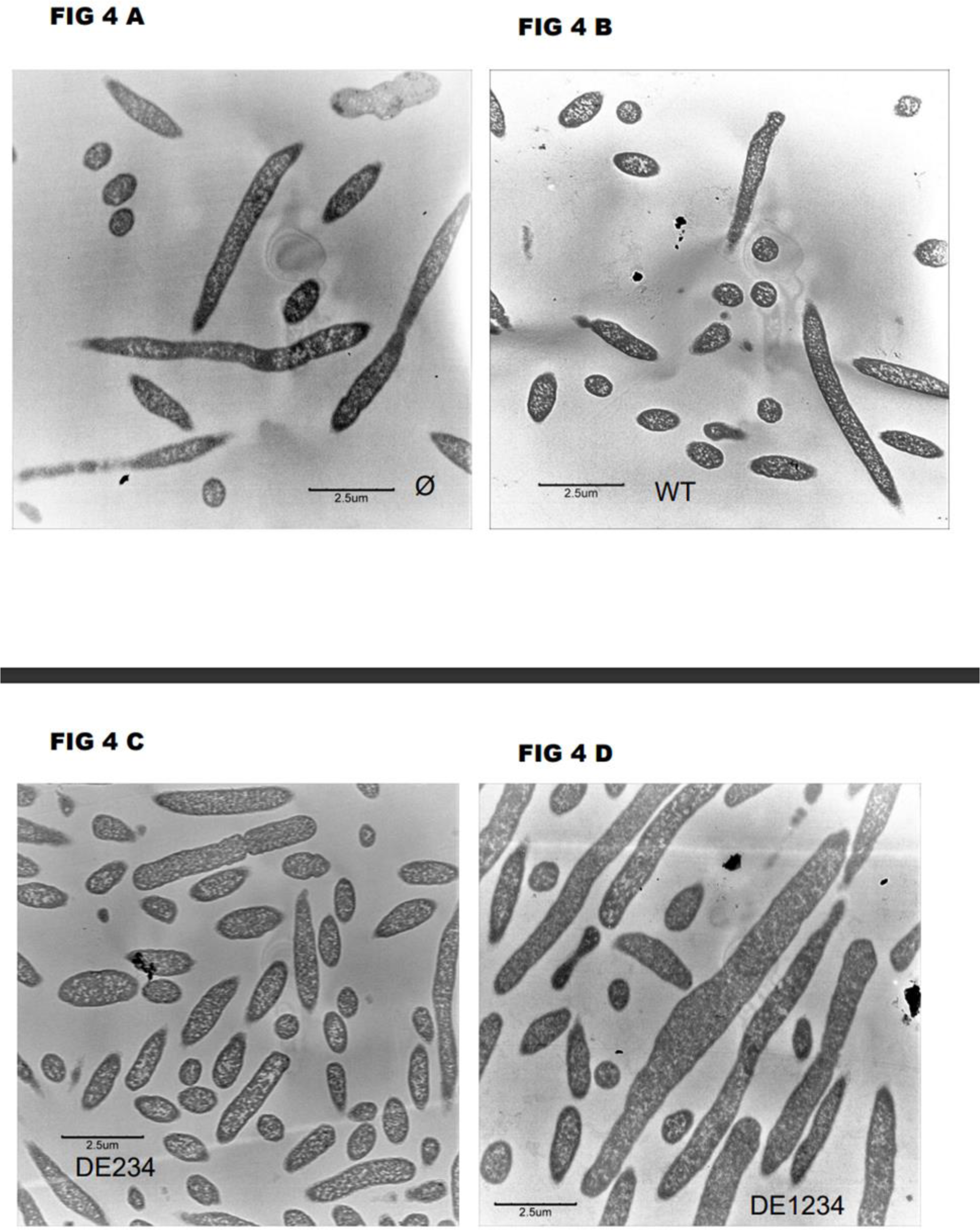
Electron micrographs of *E. coli* grown in AMPICILLIN (200 µg/ml) PLUS PIPERACILLIN (4 µg/ml) and expressing wild type or mutant annexin A4. **4A:** Ø control plasmid with no insert **4B**: WT wild type annexin A4 **3C:** DE234 mutant annexin A4 **3D:** DE1234 mutant annexin A4

## DISCUSSION

This study reports the unexpected finding that a eukaryotic calcium- and membrane-binding protein, annexin A4, can enhance cytokinesis in bacteria in which cytokinesis is partially inhibited by beta-lactam antibiotics. The processes of cytokinesis in eukaryotes and in prokaryotes have in common characteristic morphological steps. The dividing cells pinch down into a neck between the forming daughter cells, the membranes in the neck fuse together and the cells separate. The molecular events underlying these morphological changes are under intense study in both cell types [9,22–24]. Different, unrelated sets of proteins undergo complex interactions in each of these types of organisms to mediate cytokinesis, but there is little similarity or known homology between the sets of proteins. In prokaryotes like *E. coli* a master protein appears to be FtsZ which has some features in common with the eukaryotic protein tubulin in that it has a GTPase activity and undergoes polymerization [25]. The FtsZ polymer forms a ring around the middle of the bacterium that contracts to form the neck between the dividing cells leading to membrane fusion and separation of the cells. However, to accomplish this the protein interacts with numerous other proteins, some of which, like penicillin binding protein 3 (PB3), bind beta-lactam antibiotics. The set of proteins responsible for cytokinesis in eukaryotes is also complex and the complete sequence of protein interactions is not fully worked out. Beta-lactam antibiotics do not interact with the proteins participating in cytokinesis in eukaryotic cells, indicating a significant contrast between the mechanisms in the two types of cells. These differences raise the question of how a single protein, annexin A4, can influence cytokinesis processes involving such distinct protein machineries. The answer may be that the annexin interacts with two critical elements common to both types of cells: intracellular calcium and phospholipids [6]. The importance of the interaction with calcium in bacteria is demonstrated in this report since disabling the calcium binding sites in the annexin reduces its ability to promote cytokinesis. The interaction of the annexin with calcium is intimately linked to its interactions with phospholipids since both of these ligands participate in forming the high affinity calcium and phospholipid binding site in the annexin[8]. Since the binding of calcium and phospholipid has been demonstrated in the simplest model systems, involving only an annexin, calcium, and lipid bilayers, to lead to bilayer fusion[26], no other proteins may be necessary to support membrane fusion in cells. However, evidently other proteins are important in both eukaryotes and prokaryotes for membrane fusion to be utilized in the biological context of cytokinesis.

On the other hand, annexins are not absolutely required for cytokinesis in prokaryotes as demonstrated by the absence of annexins in the majority of bacteria [6]. A few eukaryotes, such as the yeast *S. cerevisiae*, can also carry out cytokinesis without depending on annexin homologues since they are not expressed in this yeast . Other mechanisms must be able to substitute for the role of the annexins in these counterexamples.

Although the exact mechanism by which the annexin promotes cytokinesis in bacteria is not clear, this should not prevent consideration of possible biotechnology applications of bacterial expression of annexins. As discussed in the INTRODUCTION, disassembly of bacterial filaments formed in the presence of antibiotics may promote the effectiveness of antibiotic chemotherapy by enabling cells of the immune system to destroy the non-filamentous forms of the bacteria [1, 16]. Among the proteins studied here, the DE234 mutant annexin A4 might be the most effective therapeutic because of its efficacy in promoting cytokinesis in the bacterial filaments.

The possibility of using annexins to suppress the deleterious effects of pathogenic bacteria will be dependent on the ability to express the eukaryotic annexins in the bacteria while they are invading a host. Multiple approaches may be available, including transfection by expression plasmids as used in this study, or introduction through phage-based transfer of genetic material [27].

## ACKNOWLEDGEMENTS

Hitoshi Sohma (Sapporo Medical University) kindly provided replacement samples of the DE234 and DE1234 annexin A4 mutants after samples in the University of Virginia collection were lost. Thurl Harris and Michael Nelson (University of Virginia) provided helpful advice and discussions.

Stacey Criswell, Natalia Dworak, and Helen Park of the Advanced Microscopy Facility, University of Virginia School of Medicine, provided technical assistance for the electron microscopy.

## FUNDING

Financial support was provided by the Office of the Vice President for Research, University of Virginia, and the Department of Pharmacology, University of Virginia.

## COMPETING INTERESTS

The University of Virginia has filed a preliminary patent application for the use of annexins as an adjuvant in antimicrobial chemotherapy.

